# Plant-TFClass: a structural classification for plant transcription factors

**DOI:** 10.1101/2022.11.22.517060

**Authors:** Romain Blanc-Mathieu, Renaud Dumas, Laura Turchi, Jérémy Lucas, François Parcy

## Abstract

Transcription factors (TFs) bind DNA at specific sequences to regulate gene expression. This universal process is achieved thanks to the DNA-binding domain (DBD) present in each TF. DBDs show a vast diversity of protein folds within and across organisms, ranging from simple long basic alpha helices to complex structural combinations of alpha, beta and loop folds. In mammals, the structural conformation of the DBDs and the way it establishes contact with DNA has been used to organize TFs in a hierarchical classification named TFClass. However, such classification is missing from plants that possess many DBD types absent from mammals. Here, we reviewed the numerous TF DBD 3D-structures and models available for plants to organize all plant TFs types following the TFClass hierarchy (Superclass/Class/Family/Subfamily). We classified most of the 55 recognized plant TF types within the existing TFClass framework. This extended classification led us to add six new classes and 34 new families corresponding to TF DBD structures absent in mammals. Plant-TFClass provides a unique resource for TF and TF binding sites comparison across TF families and across organisms.

## Introduction

Transcription factors (TFs) play essential roles in most processes occurring in living organisms. Thanks to a dedicated DNA binding domain (DBD), these proteins bind DNA at specific sequences (called TF binding sites or TFBS) and regulate the expression of associated genes. Plant genomes are known to contain a high proportion of TF-encoding genes including a collection of plant-specific TFs [1,2] that evolved during their history. The initial analysis of Arabidopsis genome identified 1500 TFs corresponding to 28 gene families of which 45% (16 families) were not found in mammals [3]. Since then, comparative genomic studies identified additional TFs and putative TF families in plant genomes [4] and various plant TFs classifications have been proposed, based on the nature of the DBD and other conserved protein domains [4–7]. With the increasing number of 3D DBD structures being solved or reliably predicted, it now becomes possible to propose a hierarchical classification for plant TFs based on the 3D structure and the DNA binding mode of their DBD. In human and mammals, TFClass is the reference structural classification [8,9]. This hierarchical classification starts with nine structurally defined superclasses, further decomposed into classes, families and subfamilies. It provides a framework for meta-analyses of TF properties and evolution [10], it facilitates comparisons between organisms and it is the reference classification used by JASPAR, the open access database for TFBS models [11].

Here we built upon the TFClass framework to classify all plant TFs types described to date. The new framework is referred to as “Plant-TFClass” (Figure 1, Table S1). For this, we reviewed experimental and predicted data on DBD structures. For each TF type, we proposed a classification when sufficient evidence exist and we left the remaining ones in a tenth ‘Yet undefined DNA-binding domains’ superclass as done in TFClass. As compared to TFClass, Plant-TFClass has additional TFs not present in mammals. The new plant TFs are of two natures: 1) TFs with a DBD absent from mammals. This was the case for six new classes (SBP, LEAFY, EIL, AP2/EREB, RHH, B3) and we explain on which basis these news classes were organized under a given superclass. Such TFs may have evolved (i) de novo, (ii) from existing DBD by rapid sequence evolution or (iii) from transposon proteins [12,13]. 2) Another type of plant-specific TFs corresponds to those possessing a DBD fold that exists in mammals but has diverged extensively, in particular with the acquisition of auxiliary domains. This usually defines new families within existing classes (e.g. HD-ZIP, GARP, MADS Type I, AHL).

**Figure 1:**
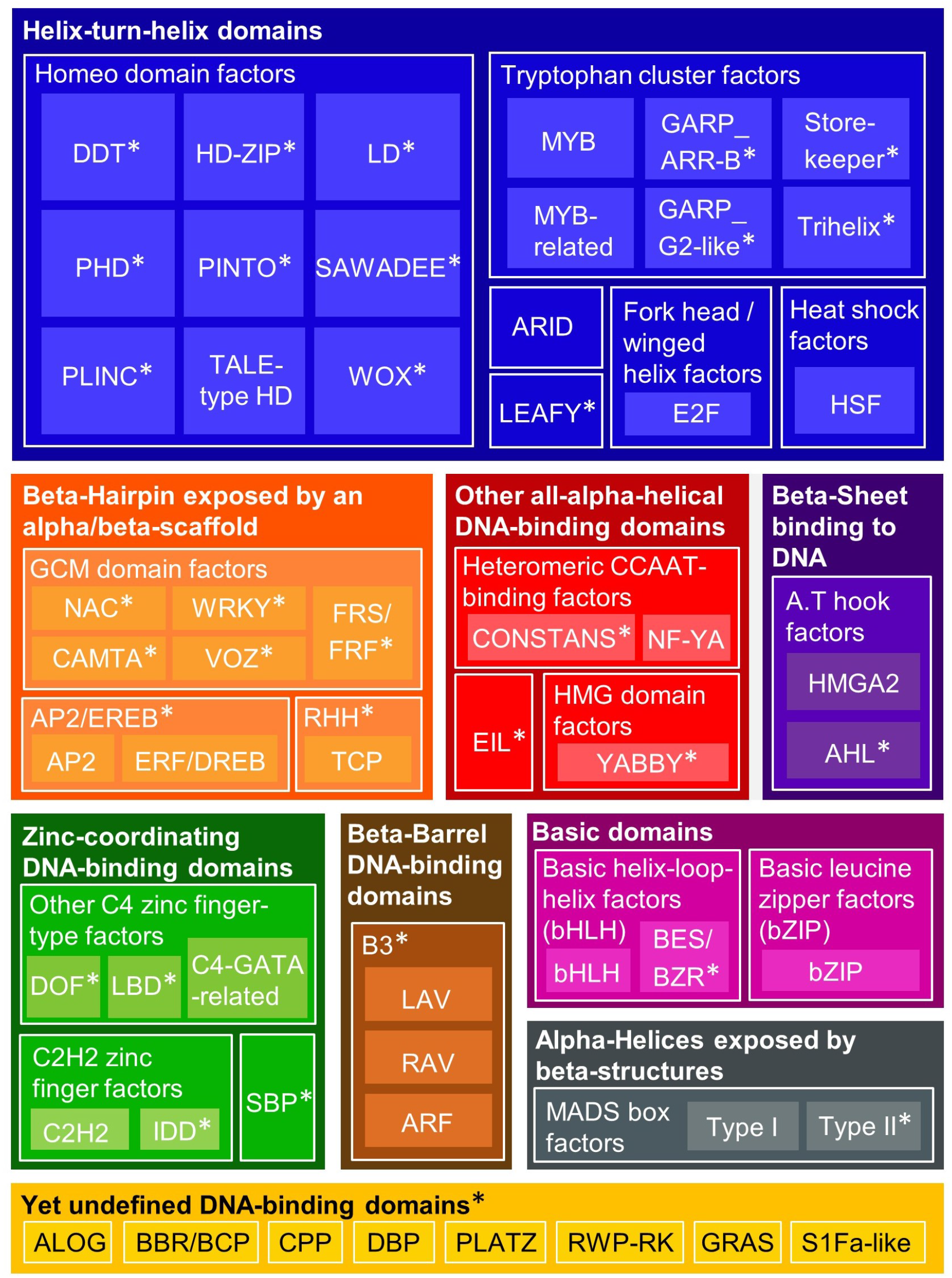
Color map illustrating the Plant-TFClass structural classification. Colors separate the nine structurally defined superclasses of transcription factors DNA binding domain plus the “Yet undefined DNA-binding domains”. Shades of color separate classes (framed with white thin lines) and families within each superclass. Asterisks indicate TF families and classes absent from TFClass. Asterisks in classes correspond to DBD fold absent in mammals.

Among 55 recognized plants TF types, 47 were classified in one of the nine superclasses from TFClass (Basic domains, Zinc-coordinating DNA-binding domains, Helix-turn-helix domains, Other all-alpha-helical DNA-binding structures, alpha-Helices exposed by beta-structures, beta-Hairpin exposed by an alpha/beta-scaffold, Immunoglobulin-fold, beta-sheet binding to DNA, beta-Barrel DNA-binding domains) and the remaining eight were added to the “Yet undefined DBD”.

### Basic domain superclass

This superclass includes TFs that bind DNA with a basic alpha-helix usually inserted in the DNA major groove. It contains two classes: the basic Leucine Zipper (bZIP) and basic Helix-Loop-Helix (bHLH) TFs that are present in most eukaryotes. Both types are dimeric TFs and pinch DNA between two basic helices. We have divided the bHLH class into two families: the classical bHLH and the BES/BZR plant-specific family. BES/BZR TFs slightly differs from classical bHLH. Indeed, instead of having only the long basic helix interacting with DNA as in classical bHLH, in BES/BZR DBDs, the loop between the two helices adds contacts with the minor groove and the second helix involved in dimerization is much shorter.

### Zinc-coordinating DNA binding domains

DBDs of TFs from this superclass have a fold organized by one or more zinc ions. The nature of the Zn coordination and the resulting DBD fold define nine classes in TFClass [9]. Within this superclass, plants possess two TFs families shared with animals: C2H2 (from the “C2H2 Zinc finger factors” class) and the C4-GATA-related (from the “Other C4 zinc finger-type factors” class). In the C2H2 fold, the “zinc finger” is a loop formed between a beta-hairpin and an alpha-helix and the Zn atom is coordinated by two cysteine and two histidine residues. Several Zn fingers can wrap around the major groove of the DNA via interactions of the alpha-helices with the major groove, as shown in Fig. 2. In plants, the IDD TFs bind DNA thanks to two C2H2 and two C2CH zinc fingers with the first C2H2 zinc finger being the most critical for specific binding [14]. We thus classified IDD TFs as a family within the C2H2 class. The C4-GATA-related fold is a variant of the C2H2 fold in which the coordination involves four cysteine residues. Available experimental structures led us to add three types of plant TFs within this superclass: the Squamosa promoter Binding Proteins (SBP), the DNA-binding One Zinc Finger (DOF) and LATERAL ORGAN BOUNDARIES (LOB) DOMAIN (LBD) proteins. SBP TFs possess two zinc-coordinating domains, with the N-terminal one being the best candidate to contact DNA according to NMR experiments (performed in the presence or absence of DNA) and DNA docking on the NMR structure (1UL4) [15]. SBP DBD is dissimilar to other known zinc-binding structures (no significant 3D alignments) thereby defining a new class in Plant-TFClass. DOF TFs bind DNA via their highly conserved DOF domain. The DOF domain forms a single zinc-finger motif of C2C2-type and was thus added to the “Other C4 zinc finger-type factors” class. The sequence-specific binding to DNA is supported by several *in vitro* functional studies [16–18] and DBD/DNA structure modelling studies [19,20]. The DOF domain is also able to form protein-protein interactions with other TFs [16,21–23]. LBD TFs bind DNA thanks to their dimeric and highly conserved LOB DBD. 3D structural analysis of the LOB domain from the wheat Ramosa2 protein (TtRa2LD, 5LY0) strongly supports that a C4 zinc finger motif is responsible for DNA binding while homodimerization is achieved via a C-terminal leucine zipper-like motif [24]. We thus classified LBD TFs as a new family in the existing Other C4 zinc finger-type factors class.

**Figure 2:**
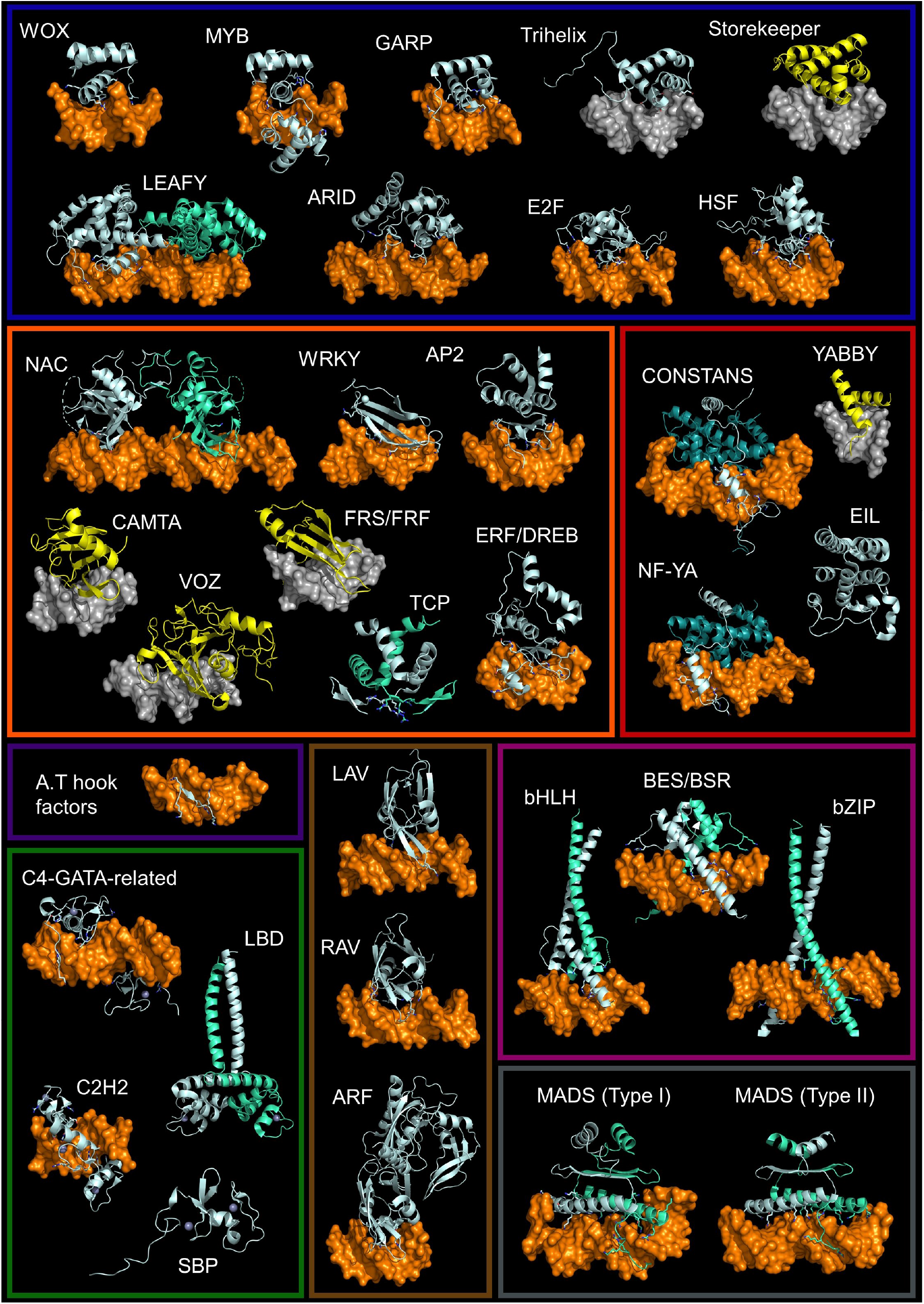
Illustration of plant TFs DNA binding domains with or without DNA. TFs are classified in their respective superclasses as in figure 1 (frames follow the superclasses color codes from Figure 1). Experimentally determined structures are colored in pale cyan (first monomer) and green cyan (second monomer). The side chains of DNA interacting residues are shown as sticks. DNA is colored orange when it was crystalized with the TF DBD and grey when it comes from another experimentally derived TF/DBD structure used in superimposition (example: the Trihelix superimposed on a MYB crystalized with DNA). When no experimental structure exists, the superimposed AlphaFold2 computed structure is shown in yellow. In superimposition the structure that was used as a template is hidden. For the two heteromeric CCAAT-binding factors, NF-YB and C are colored in deep teal. PyMOL session files are available as supplementary material.

Plant and animal genomes also share members of the C3H family that display a conserved CCCH-type domain. However, we did not include these proteins as they bind various molecules in addition to DNA. Moreover, DAP-seq experiments (DNA affinity purification followed by sequencing experiments) performed with six plants C3H proteins did not show any sequence specific DNA binding [18].

### Helix-turn-helix domains

TFs of the helix-turn-helix (HTH) domains superclass bind specific DNA sequences by inserting a ‘recognition’ alpha-helix into the major groove. This recognition helix is followed by a turn and, in most cases, two additional alpha-helices packed nearly perpendicularly against it. HTH DBDs also often have an additional structure (usually a loop) that projects basic residues in the DNA minor groove, also contributing to the sequence specificity. TFClass divides the HTH domains superclass into seven classes, five of which are present in plants: Heat shock factors, Fork head / winged helix factors, AT-rich interaction domain (ARID), Homeodomain (HD) and Tryptophan cluster factors.

The three former classes are represented by a single family in plants (HSF, E2F and ARID respectively). In flowering plants and moss, the ARID is sometimes associated with a HMG-box domain but models suggest that ARID-HMG proteins interact with DNA mainly thanks to their additional ARID domain [25]. Thus, we classified them within the ARID family.

As for the HD TF, their variety in plants comes from the diversity of adjacent domains including a leucine zipper in HD-Zip that affect DNA binding by allowing dimerization [26]. 3D structures of plant HD DBDs are available for WUSCHEL (WOX family) (6RYI) and revealed a canonical HD fold with loop regions connecting the three α-helices that are expanded compared to the drosophila Engrailed HD protein [27].

The Tryptophan cluster factors class contains MYB and MYB-related TFs families that are represented by few members in animals but greatly expanded in land plants (as R2R3-MYB: MYB genes containing two repeats of the MYB-type HTH domain) to form one of the largest transcription factor family [28]. A structure of plant MYB DBD in complex with DNA is available for the R2R3-MYB WEREWOLF (6KKS) and revealed a similar fold to that of animals MYB [29]. We have added to the Tryptophan cluster factors class, three plant specific families: the GARP, the Trihelix and the Storekeeper (STK) TFs (also known as GeBP). GARP TFs have a DBD distantly related to MYB. These TFs come in two subfamilies G2-like and ARR-B with structures available in complex with DNA for the G2-like TFs PHR1 (6J4R) [30] and LUX (5LXU) [31] and without DNA for the ARR-B TF ARR10 (1IRZ) [32]. The Trihelix motif was initially proposed to be distantly related to the C-MYB DBD based on sequence analysis [33] and this hypothesis was later confirmed by a solution structure of Arabidopsis GT-1 DBD (2JMW) [34]. STK TFs were included in this class based on the AlphaFold2 model of their DBD: this model shows similarity to HTH domains and 3D structure comparisons returns good alignments with members of the Trihelix family, MYB family and GARP/G2-like subfamily (with root mean square deviation (rmsd) of 3 Å on 80 equivalent positions out of 90 residues). All these families harbor conserved tryptophan residues (three for R2R3-MYB WEREWOLF and STK down to one for GARP) within the HTH domain, justifying their inclusion in this class.

Finally, we added to this HTH domain superclass the plant specific LEAFY (LFY) class. Crystal structure of LFY DBD in complex with DNA have been solved for *Arabidopsis thaliana* (2VY1, 2VY2) and *Physcomitrium patens* (4BHK) proteins, revealing the presence of an HTH motif [35] embedded within a 7-alpha-helices fold, plus a loop that adds base contacts in the DNA minor groove. LFY bind DNA as a dimer and possess a capacity to higher order assembly thanks to the presence of a SAM oligomerization domain [36].

### Other all-alpha-helical DNA-binding domains

This superclass gathers TFs possessing a DBD exclusively made of alpha helices but with no similarity with the aforementioned superclasses, neither for their structure nor for their mode of DNA-binding. It contains the heteromeric CCAAT-binding factors (also known as nuclear factor Y or NF-Y) and the High-mobility group (HMG) domain factors.

The former class is represented by a single family, present in many eukaryotes including plants. Heteromeric CCAAT-binding factors bind DNA as trimers made of NF-YB and NF-YC subunits plus either NF-YA or the plant-specific CONSTANS (CO) protein. The NF-YB/C dimer binds to DNA in a non-specific manner acting as a scaffold for the binding of NF-YA or CO which provide DNA sequence specificity. Despite possessing no sequence homology, NF-YA6 and CO show similar 3D-structures in complex with NF-YB/C and DNA (6R2V, 7CVQ respectively) with one long helix interacting with the NF-YB/C dimer and one short helix entering the DNA minor groove [37,38]. We thus added CO to the heteromeric CCAAT-binding factors despite that it does not recognize a CCAAT motif.

High-mobility group (HMG) domain factors represent a broad group of proteins present in all eukaryotes that bind DNA thanks to an HMG-box to remodel chromatin and to regulate gene transcription. In TFClass, this class contains seven families. Plants HMG-box containing proteins are less diversified than in animals and have either no demonstrated TF activity or a TF activity achieved via another DBD than the HMG-box (A.T hook or ARID domain) [39].

We classified the plant-specific YABBY proteins as a new family within the “High-mobility group (HMG) domain factors”. YABBY TFs possess two highly conserved domains that have been proposed to contribute to DNA binding: the N-terminal zinc finger domain and the C-terminal YABBY domain [40,41]. The YABBY domain has a helix-loop-helix domain that resembles the first two helices of the HMG-box and shows a high degree of conservation of amino acid residues that are necessary for DNA binding in HMG-boxes [40]. The AlphaFold2-predicted helix-loop-helix (39 amino acids long) of the YABBY domain structure superimposes very well (39 equivalent positions out of 39 residues, rmsd = 1.38 Å) with the HMG domain of the drosophila HMG-D protein (1QRV). This superimposition does not exclude a direct DNA binding by the zinc finger domain but experiments on the YABBY CRABS CLAW protein showed a more prominent role for the YABBY domain compared to the zinc finger domain for DNA binding [42].

To this superclass, we also added a new plant-specific class: EIL (standing for Ethylene-insensitive3 (EIN3)-like factors). The NMR solution structure of Arabidopsis EIL3 DBD without DNA (1WIJ) revealed a novel fold made of five alpha-helices with candidate DNA-contacting residues that remain to be confirmed [43].

### alpha-Helices exposed by beta-structures

Members of this superclass also have an all-alpha-helically folded DBD but contrary to the four first superclasses, the DNA-binding helices are exposed by a scaffold of beta-strands and do not insert in either DNA groove but are packed against the DNA double helix. This superclass contains two classes: the SAND domain factors and the MADS box factors. SAND domain factors are represented by ULTRAPETALA genes in plants and by VARL genes in green algae but a sequence-specific DNA binding activity has never been demonstrated [44,45]. The MADS box factors class contains two families in plants (Type I and Type II) with a DBD/DNA complex structure only described in animals. The DBD structure of Arabidopsis floral organ identity MADS TF SEPALLATA3 was recently obtained without DNA (7NB0) and did not reveal any major differences with metazoan proteins [46].

### beta-Hairpin exposed by an alpha/beta-scaffold

TFs from this superclass have a DBD made of an alpha/beta-structured scaffold with a beta-hairpin inserting into the DNA major groove and acting as the main DNA contacting element. In TFClass, this superclass contains the GCM domain factors class represented by a single mammal-specific family (the GCM family) to which we have added five families absent from mammals: the NAC, VOZ, CAMTA, WRKY and FRS/FRF. The determination of DBD/DNA crystallographic structures of the NAC TF AtANAC019 (3SWP) and the WRKY TF AtWRKY4 (2LEX) revealed that the DNA-interaction mode is analogous to that of GCM (1ODH) with good superimposition of the beta-strand’s amino acids contacting the DNA [47]. We deliberately chose to no longer classify WRKY TFs as Zn-coordinating because the domain coordinated by Zn ions lies outside their DBD. We included the Vascular Plant One-Zinc-Finger (VOZ) and the calmodulin binding transcription activator (CAMTA) families because the AlphaFold2 structural predictions of their DBDs align well with NAC DBD (3SWP) (rmsd of 3 Å on 116 equivalent positions out of 171 residues for VOZ and 2.66 Å on 94 equivalent positions out of 112 residues for CAMTA). These similarity levels are similar to WRKY (2LEX) to NAC (3SWP) comparison (55 equivalent positions out of 63 residues with an rmsd of 3.06 Å).

FRS/FRF factors have been linked to WRKY factors based on an iterative PSI-blast search [48]. AlphaFold2-predicted structure of the DBD of the FRS/FRF protein FAR1 revealed a fold similar to that of WRKY and 3D structures comparisons return significant alignment against AtWRKY4 (2LEX) (60 equivalent positions out of 90 residues with an rmsd of 3.27 Å). FRS/FRF factors were thus added to the GCM class.

We made two more additions to this superclass. We added the AP2/EREBP class corresponding to proteins with the plant specific AP2/ERF DBD [13]. The AP2/EREBP class was divided into two families: the AP2 and the ERF/DREB. In the AP2 family, DBD/DNA structures are available for TEM1 (7ET4) and AtERF1 (1GCC). Both structures revealed a beta-hairpin overhanging the major groove and topped by an alpha helix [49,50]. In the ERF/DREB family and as revealed by the TF/DNA structure of AtERF96 (5WX9), an N-terminal α-helix also contributes to the DNA interaction by entering the minor groove [51].

We have also added the TCP factors. Until recently, these plant-specific TFs were considered similar to bHLH [52]. However, the determination of the crystallographic structure of the DBD of *O. sativa* PCF6 (5ZKT) revealed they rather belong to Ribbon-Helix-Helix (RHH) TFs [53], a class yet never described in plants that recognize DNA motifs in the major groove thanks to an anti-parallel beta-sheet involved in dimer formation, anchored by alpha-helices [53,54]. We have thus added the RHH class and TCP family to the “beta-Hairpin exposed by an alpha/beta-scaffold” superclass.

### Immunoglobulin-fold

In TFClass, this superclass contains 16 families grouped into six classes that have no homologs with demonstrated TF activity in plants.

### beta-sheet binding to DNA

This superclass contains the A.T hook factors and TATA-binding proteins classes which bind through single extended strands or beta-sheets, respectively, preferentially in the minor groove of the DNA [9]. A.T hook motifs exist in a wide range of eukaryotic nuclear proteins [55]. In TFClass, the A.T hook factors class contains a single family, the HMGA, that bind DNA at AT-rich stretches thanks to a central Arg-Gly-Arg core that adopts an extended conformation deep in the minor groove [56,57]. In plants, HMGA proteins contains four A.T hook DNA binding motif with at least two required for efficient binding [58]. The overall structure of plant and animals HMGA proteins is quite different.

To this class we added the plant specific A.T hook motif nuclear-localized (AHL) transcription factors, not orthologous to the HMGA and possessing one or two A.T hook motifs. In addition AHL possess a plant and prokaryote conserved (PPC) domain, involved in protein-protein interactions [59,60].

### beta-Barrel DNA-binding domains

TF from this superclass bind DNA thanks to a beta barrel of variable number of beta-strands. It is represented by a single family in mammals (The Dbp family of the Cold-shock domain factors class). Distant homologs to this family in *A. thaliana* (CSP1 and CSP3) bind RNA to act as chaperone but have no known DNA binding activities.

On the other hand and as noted by Wingender et al. [9], the plant-specific B3 domain TFs have sequence specific DNA binding and belong to this superclass. B3 domains TFs are present in four families identified as LAV (LEAFY COTYLEDON2 (LEC2)-ABSCISIC ACID INSENSITIVE-3 (ABI3)-VAL), ABI/VP1-related protein (RAV), REPRODUCTIVE MERISTEM (REM), and AUXIN RESPONSE FACTOR (ARF) [61]. ARFs and LAVs have a single B3 domain, whereas REM family proteins have from one to eleven B3 domains [62]. RAVs have an AP2/ERF DNA binding domain in addition to the B3 domain. The B3 domain contains approximately 110 residues with seven β-strands and two α-helices folded into a pseudo-barrel. The residues belonging to the loops between the β-strands 1-2 and 4-5 contact bases in the DNA major groove allowing each B3 family to recognize a specific sequence [63–65]. Most ARF TFs possess a PB1 oligomerization domain [66–68] and often bind DNA as dimers with preference regarding the orientation and the spacing of ARF binding sites [18,69–71]. In contrast to the other families, studies performed on VRN1 [72] and REM16 [73] belonging to the REM family suggest that REMs do not have specific DNA binding sequences.

### Yet undefined DNA-binding domains

This superclass gathers mostly TFs with functionally well-characterized DBD but awaiting structural data for a definitive classification. In TFClass, it includes four classes, plus an “uncharacterized” class gathering six families. Each class is represented by a single family whose DBD is characterized to some extent. Among those families, plants possess NFX1-like proteins whose TF activity remains to be firmly established [74].

To this superclass, we added height classes corresponding to plant factor families with well characterized TF activity but lacking crucial structural clues on the DBD/DNA interaction. For three families (the cysteine-rich polycomb-like protein (CPP) [75], the DNA-binding protein phosphatase (DBP) [76] and the S1Fa-like factors [77]), there is no structural data and AlphaFold2 predictions of the 3D structure for their DBD were uninformative (low confidence model or no hit in 3D-structure comparison). Putative classification is discussed below for 5 families (GRAS, RWP-RK, PLATZ, ALOG and BCP) based on experimental or modeled structures.

The GRAS domain of SCARECROW-LIKE7 from *Oryza sativa* was crystallized as a dimer (5HYZ) with each monomer consisting of a core region made of alpha and beta structures, topped with an alpha helical cap structure. The two cap regions form a candidate DNA-binding groove containing positives residues and DNA docking in this groove predicted tight protein/DNA interactions that were validated experimentally [78]. Once validated at the structural level, such DNA binding mode would define a new superclass.

RWP-RK or NIN-like Protein (NLP) factors possess RWP-RK DNA binding domain with a basic region [79] predicted to fold as three short and one long alpha-helices (AlphaFold2 model from Uniprot). These plant-specific TFs have been shown to bind DNA as dimers [80]. 3D-structures comparison revealed significant alignment with the helix-turn-helix motif of bacterial DNA binding protein FIS (5DS9) (42 equivalent positions out of 82 residues with an rmsd of 3.01 Å) helices. RWP-RK could thus define a new class within the helix-turn-helix superclass.

The plant-specific PLATZ TFs are zinc-dependent DNA binding proteins. The zinc finger coordination motif has, in its N-terminal part, a consensus signature C-x2-H-x11-C-x2-C-x(4– 5)-C-x2-C-x(3–7)-H-x2-H which is different from other characterized zinc-binding motifs. It also has four conserved cysteine residues in its central region [81]. Structural models show various folds (alpha helices, a beta-hairpin and a beta-sheet). In the absence of experimental structure in complex with DNA or extensive modeling, it is not possible to firmly place this family within the Zinc-coordinating DNA-binding domains superclass.

With functional roles established in Arabidopsis, rice, tomato and *Marchantia polymorpha,*the ALOG (Arabidopsis LSH1 and Oryza G1) family of proteins was proposed as a novel plant specific TF family but is still poorly characterized at the structural level [82]. These proteins likely originated from a recombinase of a retrotransposon in which a Zinc ribbon was inserted [83] into a four-alpha-helix bundle DBD. Once the DNA binding specificity and the structure of the DBD in complex with DNA are characterized, these putative TFs might be classified either in the basic domain superclass or in the Zinc coordinating DNA-binding proteins.

The BCP DBD has a conserved WAR/KHGTN motif required for DNA binding, reminiscent of the WRKYCGK consensus of WRKY proteins suggesting that it may bind DNA in similar way [84].

## Method

Experimentally-determined 3D structures were downloaded from the Protein Data Bank. AlphaFold2 [85] computed structure models were downloaded from UniProt. For each model, position of the DBD was obtained directly from the UniProt “Family and Domains” annotation or identified using CD-search against the CDD database [86]. Structure predictions at the DBD positions were extracted from the PDB files using the Gemmi library (https://gemmi.readthedocs.io/en/latest/). Protein 3D structure comparisons and representation were performed using DALI [87], FATCAT [88] and PyMOL. In the text and for the sake of clarity, we colored names of TF classes and families in blue and orange, respectively.

## Supporting information

PyMOL session files

## Supplementary material

The Plant-TFClass classification is provided in a tabular format in supplemental table 1. Structure representations in figure 2 are available as PyMOL session files.

## Author contributions

FP conceived the study. RBM and RD defined the plant structural classes. All authors performed the classification. RBM, FP and RD wrote the manuscript.

## Acknowledgments

We thanks Edgar Wingender, Chloe Zubieta and Gabrielle Tichtinsky for their advices on this work. This work was supported by the French National Research Agency in the framework of the “Investissements d’avenir” program (ANR-15-IDEX-02) to RBM, by the ANR-18-CE12-0014 ChromAuxi project to RD and by the ANR-17-CE20-0014-01 Ubiflor and ANR-21-CE20-0024 Beflore projects to FP, and a PhD Fellowship from CNRS Prime80 to LT.

## References

1. Shiu, S.-H. et al. (2005) Transcription Factor Families Have Much Higher Expansion Rates in Plants than in Animals. Plant Physiol 139, 18–26

2. De Mendoza, A. et al. (2013) Transcription factor evolution in eukaryotes and the assembly of the regulatory toolkit in multicellular lineages. Proceedings of the National Academy of Sciences of the United States of America 110, E4858–E4866

3. Riechmann, J.L. et al. (2000) Arabidopsis transcription factors: genome-wide comparative analysis among eukaryotes. Science 290, 2105–2110

4. Wilhelmsson, P.K.I. et al. (2017) Comprehensive Genome-Wide Classification Reveals That Many Plant-Specific Transcription Factors Evolved in Streptophyte Algae. Genome Biology and Evolution 9, 3384–3397

5. Mukherjee, K. et al. (2009) A Comprehensive Classification and Evolutionary Analysis of Plant Homeobox Genes. Mol Biol Evol 26, 2775–2794

6. Jin, J. et al. (2017) PlantTFDB 4.0: toward a central hub for transcription factors and regulatory interactions in plants. Nucleic Acids Research 45, D1040–D1045

7. Yilmaz, A. et al. (2011) AGRIS: the Arabidopsis Gene Regulatory Information Server, an update. Nucleic Acids Research 39, D1118–D1122

8. Wingender, E. et al. (2018) TFClass: expanding the classification of human transcription factors to their mammalian orthologs. Nucleic Acids Res 46, D343–D347

9. Wingender, E. (2013) Criteria for an updated classification of human transcription factor DNA-binding domains. J Bioinform Comput Biol 11, 1340007

10. Ambrosini, G. et al. (2020) Insights gained from a comprehensive all-against-all transcription factor binding motif benchmarking study. Genome Biology 21, 114

11. Castro-Mondragon, J.A. et al. (2022) JASPAR 2022: the 9th release of the open-access database of transcription factor binding profiles. Nucleic Acids Research 50, D165–D173

12. de Mendoza, A. and Sebé-Pedrós, A. (2019) Origin and evolution of eukaryotic transcription factors. Current Opinion in Genetics & Development 58–59, 25–32

13. Yamasaki, K. et al. (2013) DNA-binding domains of plant-specific transcription factors: structure, function, and evolution. Trends in Plant Science 18, 267–276

14. Hirano, Y. et al. (2017) Structure of the SHR-SCR heterodimer bound to the BIRD/IDD transcriptional factor JKD. Nat Plants 3, 17010

15. Yamasaki, K. et al. (2004) A novel zinc-binding motif revealed by solution structures of DNA-binding domains of Arabidopsis SBP-family transcription factors. J Mol Biol 337, 49–63

16. Yanagisawa, S. (2004) Dof Domain Proteins: Plant-Specific Transcription Factors Associated with Diverse Phenomena Unique to Plants. Plant and Cell Physiology 45, 386–391

17. Umemura, Y. et al. (2004) The Dof domain, a zinc finger DNA-binding domain conserved only in higher plants, truly functions as a Cys2/Cys2 Zn finger domain. The Plant Journal 37, 741–749

18. O’Malley, R.C. et al. (2016) Cistrome and Epicistrome Features Shape the Regulatory DNA Landscape. Cell 165, 1280–1292

19. Hamzeh-Mivehroud, M. et al. (2015) Identifying key interactions stabilizing DOF zinc finger–DNA complexes using in silico approaches. Journal of Theoretical Biology 382, 150–159

20. Pandey, B. et al. (2018) Dynamics of Dof domain-DNA interaction in wheat: Insights from atomistic simulations and free energy landscape. Journal of Cellular Biochemistry 119, 8818–8829

21. Zhang, B. et al. (1995) Interactions between distinct types of DNA binding proteins enhance binding to ocs element promoter sequences. Plant Cell 7, 2241–2252

22. Ruta, V. et al. (2020) The DOF Transcription Factors in Seed and Seedling Development. Plants (Basel) 9, 218

23. Gao, H. et al. (2021) PIF4 enhances DNA binding of CDF2 to co-regulate target gene expression and promote Arabidopsis hypocotyl cell elongation. Nat. Plants 8, 1082–1093

24. Chen, W.-F. et al. (2019) Structural analysis reveals a “molecular calipers” mechanism for a LATERAL ORGAN BOUNDARIES DOMAIN transcription factor protein from wheat. J Biol Chem 294, 142–156

25. Roy, A. et al. (2016) Deciphering the role of the AT-rich interaction domain and the HMG-box domain of ARID-HMG proteins of Arabidopsis thaliana. Plant Mol Biol 92, 371–388

26. Bürglin, T.R. and Affolter, M. (2016) Homeodomain proteins: an update. Chromosoma 125, 497–521

27. Sloan, J. et al. (2020) Structural basis for the complex DNA binding behavior of the plant stem cell regulator WUSCHEL. Nat Commun 11, 2223

28. Du, H. et al. (2015) The Evolutionary History of R2R3-MYB Proteins Across 50 Eukaryotes: New Insights Into Subfamily Classification and Expansion. Sci Rep 5, 11037

29. Wang, B. et al. (2020) Structural insights into target DNA recognition by R2R3-MYB transcription factors. Nucleic Acids Research 48, 460–471

30. Jiang, M. et al. (2019) Structural basis for the Target DNA recognition and binding by the MYB domain of phosphate starvation response 1. FEBS J 286, 2809–2821

31. Silva, C.S. et al. (2016) The Myb domain of LUX ARRHYTHMO in complex with DNA: expression, purification and crystallization. Acta Crystallogr F Struct Biol Commun 72, 356–361

32. Hosoda, K. et al. (2002) Molecular Structure of the GARP Family of Plant Myb-Related DNA Binding Motifs of the Arabidopsis Response Regulators. Plant Cell 14, 2015–2029

33. Nagano, Y. (2000) Several features of the GT-factor trihelix domain resemble those of the Myb DNA-binding domain. Plant Physiol 124, 491–494

34. Nagata, T. et al. (2010) Solution structures of the trihelix DNA-binding domains of the wild-type and a phosphomimetic mutant of Arabidopsis GT-1: Mechanism for an increase in DNA-binding affinity through phosphorylation. Proteins: Structure, Function, and Bioinformatics 78, 3033–3047

35. Hamès, C. et al. (2008) Structural basis for LEAFY floral switch function and similarity with helix-turn-helix proteins. The EMBO Journal 27, 2628–2637

36. Sayou, C. et al. (2016) A SAM oligomerization domain shapes the genomic binding landscape of the LEAFY transcription factor. Nature Communications 7, 11222

37. Chaves-Sanjuan, A. et al. (2021) Structural determinants for NF-Y subunit organization and NF-Y/DNA association in plants. The Plant Journal 105, 49–61

38. Lv, X. et al. (2021) Structural insights into the multivalent binding of the Arabidopsis FLOWERING LOCUS T promoter by the CO–NF–Y master transcription factor complex. The Plant Cell 33, 1182–1195

39. Antosch, M. et al. (2012) Plant Proteins Containing High Mobility Group Box DNA-Binding Domains Modulate Different Nuclear Processes1[W]. Plant Physiol 159, 875–883

40. Bowman, J.L. and Smyth, D.R. (1999) CRABS CLAW, a gene that regulates carpel and nectary development in Arabidopsis, encodes a novel protein with zinc finger and helix-loop-helix domains. Development 126, 2387–2396

41. Sawa, S. et al. (1999) FILAMENTOUS FLOWER, a meristem and organ identity gene of Arabidopsis, encodes a protein with a zinc finger and HMG-related domains. Genes Dev 13, 1079–1088

42. Gross, T. et al. (2018) CRABS CLAW Acts as a Bifunctional Transcription Factor in Flower Development. Frontiers in Plant Science 9

43. Yamasaki, K. et al. (2005) Solution Structure of the Major DNA-binding Domain of Arabidopsis thaliana Ethylene-insensitive3-like3. Journal of Molecular Biology 348, 253–264

44. Carles, C.C. and Fletcher, J.C. (2009) The SAND domain protein ULTRAPETALA1 acts as a trithorax group factor to regulate cell fate in plants. Genes Dev. 23, 2723–2728

45. Duncan, L. et al. (2006) Orthologs and paralogs of regA, a master cell-type regulatory gene in Volvox carteri. Curr Genet 50, 61–72

46. Lai, X. et al. (2021) The intervening domain is required for DNA-binding and functional identity of plant MADS transcription factors. Nature Communications 12

47. Welner, D.H. et al. (2012) DNA binding by the plant-specific NAC transcription factors in crystal and solution: a firm link to WRKY and GCM transcription factors. Biochem J 444, 395–404

48. Babu, M.M. et al. (2006) The natural history of the WRKY–GCM1 zinc fingers and the relationship between transcription factors and transposons. Nucleic Acids Research 34, 6505–6520

49. Allen, M.D. et al. (1998) A novel mode of DNA recognition by a beta-sheet revealed by the solution structure of the GCC-box binding domain in complex with DNA. EMBO J 17, 5484–5496

50. Hu, H. et al. (2021) TEM1 combinatorially binds to FLOWERING LOCUS T and recruits a Polycomb factor to repress the floral transition in Arabidopsis. Proc Natl Acad Sci U S A 118,e2103895118

51. Chen, C.-Y. et al. (2020) Structural insights into Arabidopsis ethylene response factor 96 with an extended N-terminal binding to GCC box. Plant Mol Biol 104, 483–498

52. Kosugi, S. and Ohashi, Y. (1997) PCF1 and PCF2 specifically bind to cis elements in the rice proliferating cell nuclear antigen gene. Plant Cell 9, 1607–1619

53. Sun, L. et al. (2020) The crystal structure of the TCP domain of PCF6 in Oryza sativa L. reveals an RHH-like fold. FEBS Letters 594, 1296–1306

54. Schreiter, E.R. and Drennan, C.L. (2007) Ribbon–helix–helix transcription factors: variations on a theme. Nat Rev Microbiol 5, 710–720

55. Aravind, L. and Landsman, D. (1998) AT-hook motifs identified in a wide variety of DNA-binding proteins. Nucleic Acids Res 26, 4413–4421

56. Fonfría-Subirós, E. et al. (2012) Crystal structure of a complex of DNA with one AT-hook of HMGA1. PLoS One 7, e37120

57. Huth, J.R. et al. (1997) The solution structure of an HMG-I(Y)–DNA complex defines a new architectural minor groove binding motif. Nat Struct Mol Biol 4, 657–665

58. Grasser, K.D. (2003) Chromatin-associated HMGA and HMGB proteins: versatile co-regulators of DNA-dependent processes. Plant Mol Biol 53, 281–295

59. Yun, J. et al. (2012) The AT-hook Motif-containing Protein AHL22 Regulates Flowering Initiation by Modifying FLOWERING LOCUS T Chromatin in Arabidopsis. J Biol Chem 287, 15307–15316

60. Meijer, A.H. et al. (1996) Novel members of a family of AT hook-containing DNA-binding proteins from rice are identified through their in vitro interaction with consensus target sites of plant and animal homeodomain proteins. Plant Mol Biol 31, 607–618

61. Swaminathan, K. et al. (2008) The plant B3 superfamily. Trends in Plant Science 13, 647–655

62. Romanel, E.A.C. et al. (2009) Evolution of the B3 DNA Binding Superfamily: New Insights into REM Family Gene Diversification. PLOS ONE 4, e5791

63. Ulmasov, T. et al. (1997) ARF1, a transcription factor that binds to auxin response elements. Science 276, 1865–1868

64. Kagaya, Y. et al. (1999) RAV1, a novel DNA-binding protein, binds to bipartite recognition sequence through two distinct DNA-binding domains uniquely found in higher plants. Nucleic Acids Research 27, 470–478

65. Mönke, G. et al. (2004) Seed-specific transcription factors ABI3 and FUS3: molecular interaction with DNA. Planta 219, 158–166

66. Korasick, D.A. et al. (2014) Molecular basis for AUXIN RESPONSE FACTOR protein interaction and the control of auxin response repression. Proceedings of the National Academy of Sciences 111, 5427–5432

67. Nanao, M.H. et al. (2014) Structural basis for oligomerization of auxin transcriptional regulators. Nat Commun 5, 3617

68. Han, M. et al. (2014) Structural basis for the auxin-induced transcriptional regulation by Aux/IAA17. Proc Natl Acad Sci U S A 111, 18613–18618

69. Boer, D.R. et al. (2014) Structural basis for DNA binding specificity by the auxin-dependent ARF transcription factors. Cell 156, 577–589

70. Stigliani, A. et al. (2019) Capturing Auxin Response Factors Syntax Using DNA Binding Models. Molecular Plant 12, 822–832

71. Cancé, C. et al. (2022) Auxin response factors are keys to the many auxin doors. New Phytologist 235, 402–419

72. Levy, Y.Y. et al. (2002) Multiple Roles of Arabidopsis VRN1 in Vernalization and Flowering Time Control. Science 297, 243–246

73. Yu, Y. et al. (2020) Arabidopsis REM16 acts as a B3 domain transcription factor to promote flowering time via directly binding to the promoters of SOC1 and FT. The Plant Journal 103, 1386–1398

74. Lisso, J. et al. (2012) NFXL2 modifies cuticle properties in Arabidopsis. Plant Signal Behav 7, 551–555

75. Cvitanich, C. et al. (2000) CPP1, a DNA-binding protein involved in the expression of a soybean leghemoglobin c3 gene. Proc Natl Acad Sci U S A 97, 8163–8168

76. Carrasco, J.L. et al. (2005) A novel DNA-binding motif, hallmark of a new family of plant transcription factors. Plant Physiol 137, 602–606

77. Santi, L. et al. (2003) The GA octodinucleotide repeat binding factor BBR participates in the transcriptional regulation of the homeobox gene Bkn3. Plant J 34, 813–826

78. Li, S. et al. (2016) Crystal Structure of the GRAS Domain of SCARECROW-LIKE7 in Oryza sativa. The Plant Cell 28, 1025–1034

79. Schauser, L. et al. (1999) A plant regulator controlling development of symbiotic root nodules. Nature 402, 191–195

80. Nishida, H. et al. (2021) Different DNA-binding specificities of NLP and NIN transcription factors underlie nitrate-induced control of root nodulation. Plant Cell 33, 2340–2359

81. Nagano, Y. et al. (2001) A novel class of plant-specific zinc-dependent DNA-binding protein that binds to A/T-rich DNA sequences. Nucleic Acids Res 29, 4097–4105

82. Naramoto, S. et al. (2020) The origin and evolution of the ALOG proteins, members of a plant-specific transcription factor family, in land plants. J Plant Res 133, 323–329

83. Iyer, L.M. and Aravind, L. (2012) ALOG domains: provenance of plant homeotic and developmental regulators from the DNA-binding domain of a novel class of DIRS1-type retroposons. Biology Direct 7, 39

84. Theune, M.L. et al. (2019) Phylogenetic Analyses and GAGA-Motif Binding Studies of BBR/BPC Proteins Lend to Clues in GAGA-Motif Recognition and a Regulatory Role in Brassinosteroid Signaling. Frontiers in Plant Science 10

85. Jumper, J. et al. (2021) Highly accurate protein structure prediction with AlphaFold. Nature 596, 583–589

86. Lu, S. et al. (2020) CDD/SPARCLE: the conserved domain database in 2020. Nucleic Acids Res 48, D265–D268

87. Holm, L. (2022) Dali server: structural unification of protein families. Nucleic Acids Research 50, W210–W215

88. Li, Z. et al. (2020) FATCAT 2.0: towards a better understanding of the structural diversity of proteins. Nucleic Acids Research 48, W60–W64

